# Warm nights disrupt global transcriptional rhythms in field-grown rice panicles

**DOI:** 10.1101/702183

**Authors:** Jigar S. Desai, Lovely Mae F. Lawas, Ashlee M. Valente, Adam R. Leman, Dmitry O. Grinevich, S.V. Krishna Jagadish, Colleen J. Doherty

## Abstract

In rice, a small increase in nighttime temperatures reduces grain yield and quality. How warm nighttime temperatures (**WNT**) produce these detrimental effects is not well understood, especially in field conditions where the normal day to night temperature fluctuation exceeds the mild increase in nighttime temperature. We observed genome-wide disruption of gene expression timing during the reproductive phase on field-grown rice panicles acclimated to 2-3°C WNT. Rhythmically expressed transcripts were more sensitive to WNT than non-rhythmic transcripts. The system-wide transcriptional perturbations suggest that WNT disrupts the tight temporal coordination between internal molecular events and the environment resulting in reduced productivity. We identified transcriptional regulators whose predicted targets are enriched for sensitivity to WNT. The affected transcripts and candidate regulators identified through our network analysis explain molecular mechanisms driving sensitivity to WNT and candidates that can be targeted to enhance tolerance to WNT.

## INTRODUCTION

Global climate models predict with high certainty that mean surface temperatures will increase by 1° to 4°C by 2100^1–3^. A breakdown of these temperature trends highlights a more rapid increase in minimum nighttime temperature compared to the maximum daytime temperature at the global^1^, regional^4^, and farm^5, 6^ scales. In contrast to the short duration heat-spikes predicted with the increasing daytime temperatures, the duration of warmer nighttime temperatures (**WNT**) is expected to increase; impacting important growth and developmental phases of crops^7^. In response to increased daytime temperatures, rice plants employ mechanisms to minimize heat-induced damage such as avoidance through transpiration cooling^8^, escape through early morning flowering^9^, and reproductive resilience^10^. In contrast, there is limited plasticity in domesticated rice plants to overcome the impacts of increasing nighttime temperature^7^. The negative impacts of WNT on rice yield and quality have been documented across controlled environments^11, 12^ and field conditions^13, 14^, demonstrating the potential to induce substantial economic losses^15^. The limited physiological capacity and the larger spatial scales of the predicted increase in nighttime temperatures compared to location-specific daytime temperature increases^7^ suggest that the economic losses under current and future warmer nights pose a severe threat to sustaining global rice production.

The physiological responses in rice to high nighttime temperatures include a significant reduction in pollen viability, increased spikelet sterility, and membrane damage leading to yield losses^11, 16–18^. However, these investigations imposed temperatures that are significantly higher than future predictions. Thus, a knowledge gap exists between rice responses in controlled chambers and real-world conditions. A series of studies using field-based heat tents demonstrated the difference between chamber and field studies^7, 13, 19–21^. These field-based studies identify higher night respiration during post-flowering as a critical factor determining yield and quality losses due to high nighttime temperature^20^. Although the relationship between night respiration and sugar metabolism enzymes has been documented, particularly during the grain-filling stage^20^, the mechanistic changes are yet to be investigated.

In the field, environmental conditions change dynamically. This variation is not captured by controlled environments and only partially by the field-based heat tents. Previous observations indicate that the environmental variability of natural field conditions plays an essential role in regulating transcriptional responses and contributes to the stability of the circadian clock in rice^22^. Extensive studies in Arabidopsis have demonstrated that plant responses to abiotic stress are dynamic throughout the day^23–38^ supporting the need to capture stress responses at multiple time points to provide a comprehensive mechanistic understanding of rice exposed to WNT. Examination of the temporal mechanistic responses to stress in crops under field conditions is generally lacking and is the primary motivation driving our investigations.

The circadian clock temporally coordinates the molecular activities with the surrounding environment^39, 40^. The clock is sensitive to subtle changes in the environment to ensure an organism is ‘in-tune’ with the surroundings^41^. WNT may disrupt this environmental coordination. In rice and Arabidopsis, daily rhythms of temperature, also known as thermocycles, entrain the circadian clock and control the rhythmic expression of a portion of the transcriptome^42–44^. In Arabidopsis, the photoreceptor PHYB connects changes in ambient temperature to the circadian clock^45–47^ and changes in ambient temperatures affect the expression of the core circadian components^48^. Although these studies indicate possible mechanisms for how plants sense temperature changes, it is still not clear how the daily temperature range is perceived and integrated into the circadian clock. Moreover, the significance of a thermocycle sensitive clock, the impacts of altering the daily temperature cycles, and the effects on the expression of these temperature-responsive rhythms, particularly under field conditions, remain to be characterized. Under WNT, the daily thermocycle amplitude is reduced and could impact the expression of thermocycle-regulated transcripts.

In order to fill the significant knowledge gap on the molecular responses inducing yield losses in field conditions under WNT, we investigated how WNT, at levels in-line with the Intergovernmental Panel on Climate Change (IPCC) predictions, affect the genome-wide expression patterns in the panicles of rice grown under field conditions. Specific objectives of our study were to (i) quantify the diurnal reprogramming of the rice panicle transcriptome under WNT; (ii) identify major interacting molecular pathways that determine rice response to WNT and (iii) find regulators of transcriptionally responsive genes under WNT.

## RESULTS

### WNT NEGATIVELY IMPACT BIOMASS AND YIELD

IR64, a popular high-yielding rice variety, was grown under normal nighttime temperatures (**NNT**) or under warm nighttime temperatures (**WNT**), using a field-based infrared ceramic heating system (Fig. S1). WNT treatment started at panicle initiation and continued through maturity. At 50% flowering, field-grown rice panicles were collected for transcriptional analysis throughout the 24h cycle. WNT maintained a 2-3°C increase in temperature in the 12h night period (1800—0600h) compared to ambient temperature (Fig. S1). As previously reported^5, 14^, we observed a 12.5% decrease in the grain yield of IR64 under WNT (Kruskal-Wallis test *p*-value <0.05, Fig. 1). Total aboveground biomass (*p*-value <0.05), number of spikelets per panicle (*p*-value <0.05), and 1000-grain weight (*p*-value*<*0.05) were also significantly affected by WNT (Fig. 1, Dataset S1). Panicles per m^2^, and spikelet fertility did not change significantly with WNT (Supplemental Table 1).

**Figure 1.**
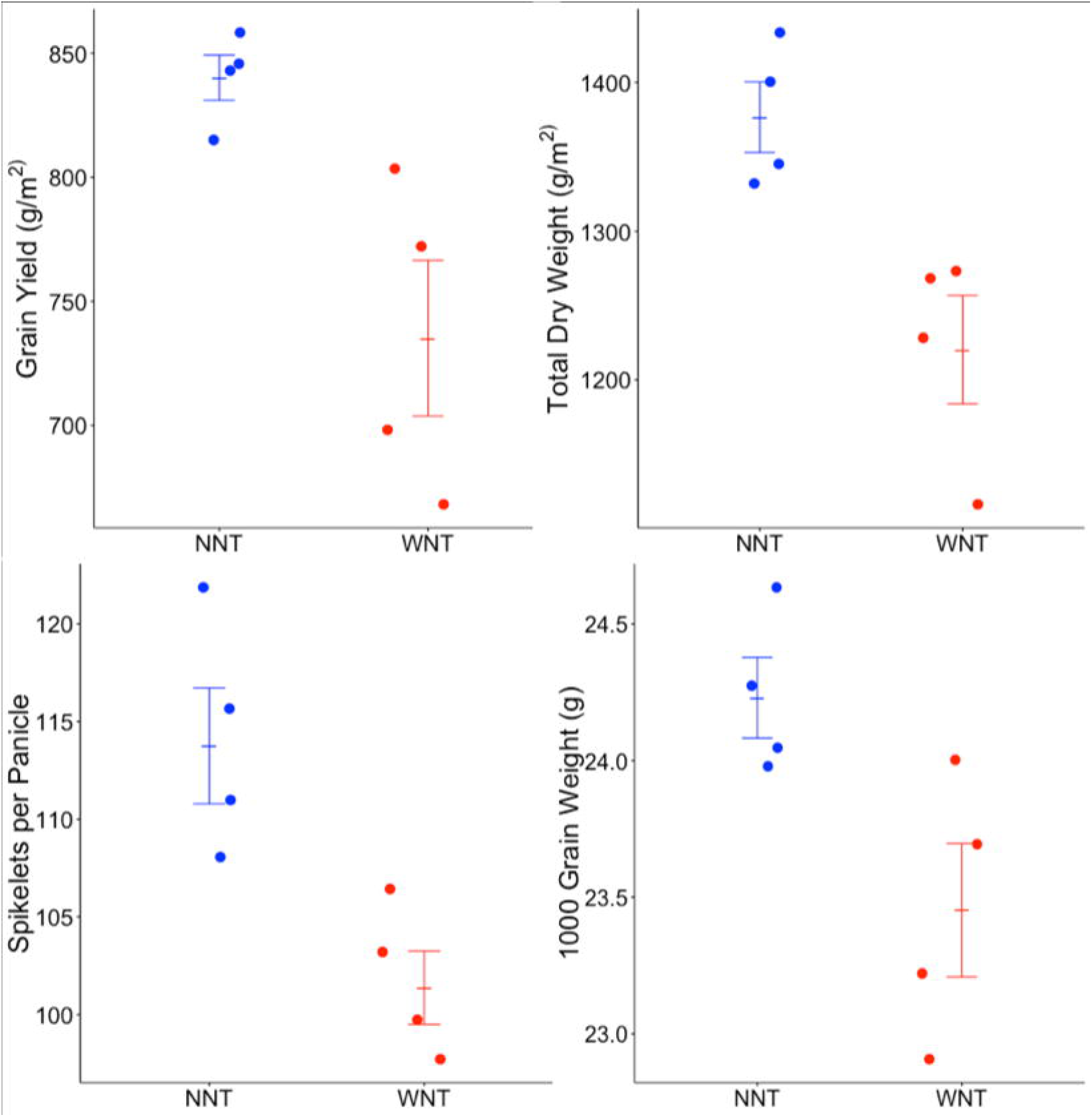
WNT impacts on agronomic performance. The effects of the Warm Nighttime Temperatures treatment (WNT, Red) compared to Normal Nighttime Temperatures (NNT, Blue) on agronomic traits of grain yield (g/m^2^), average 1000-grain weight (g), spikelets per panicle, and biomass (g/m^2^). Twelve plants were sampled from each of four plots per treatment. Error bars indicate ±SE (n=4).

### WNT IMPACTS TRANSCRIPTION PATTERNS DURING THE DAY

To evaluate the molecular changes associated with the observed agronomic changes in plants grown under WNT, samples were collected at multiple time points throughout the day. We performed RNA-Seq on panicles at the 50% flowering stage. A total of 1110 genes were identified as differentially expressed genes (DEGs) between WNT and NNT (adjusted *p*-value < 0.05 and log fold change > 0.5), corresponding to 6% of the 15,213 reliably detectable genes (Fig. 2, Supplemental Table 2). In response to WNT, 415 genes were upregulated and 695 genes were downregulated (Fig. 2A and B). The majority of the 415 DEGs that were upregulated were identified from the daytime samples, while significantly downregulated genes were more often detected in the nighttime. The expression of all detectible genes is available at https://go.ncsu.edu/clockworkviridi_wnt.

**Figure 2.**
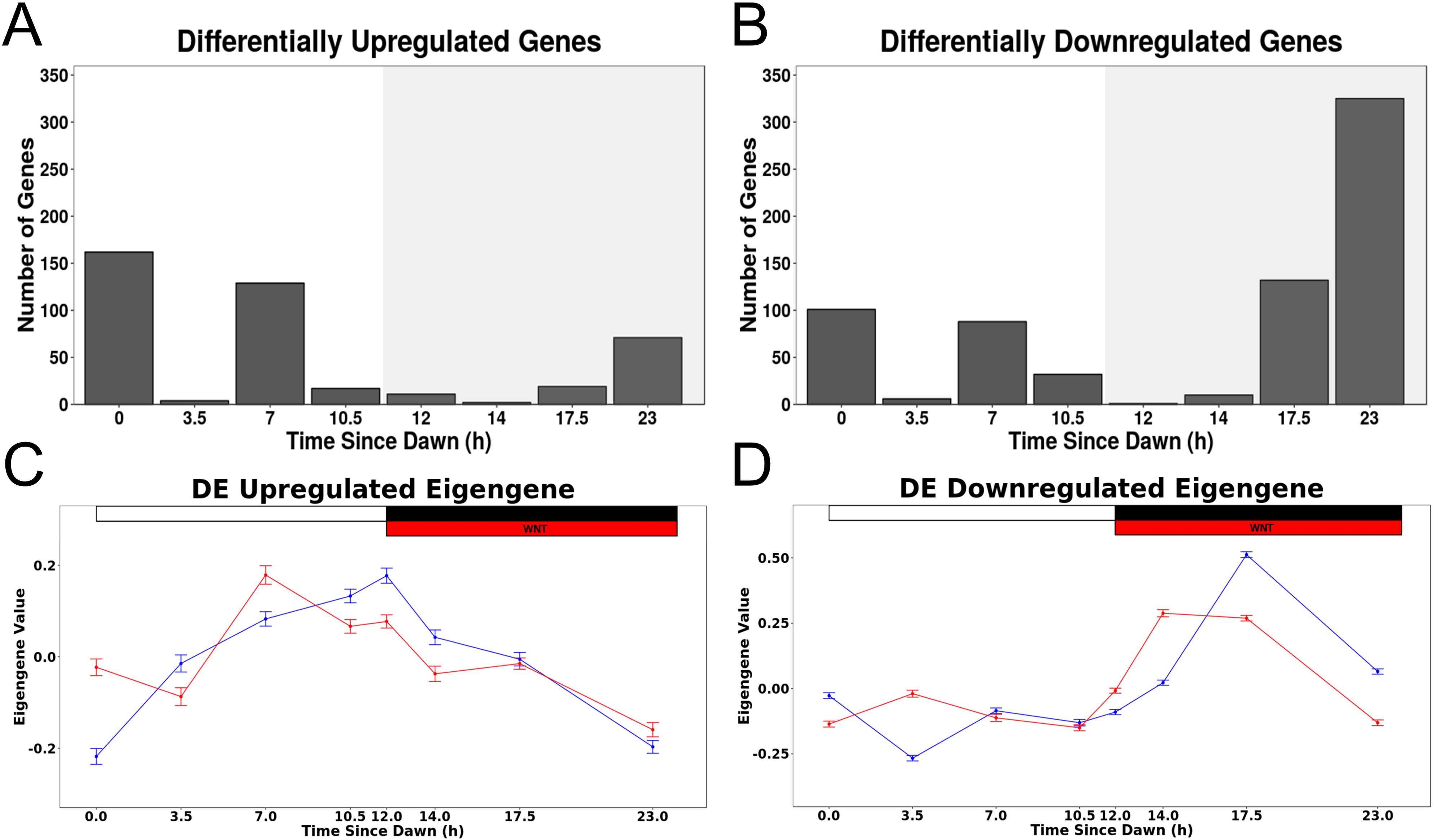
Differentially expressed genes in response to WNT. Differentially expressed genes identified by comparing expression levels between Warm Nighttime Temperatures (WNT) and Normal Nighttime Temperatures (NNT) at each time point (FDR < 0.05 and logFC > 0.5). Time points indicate sample time in hours after dawn. Number of transcripts (A) upregulated and (B) downregulated in WNT compared to NNT at each time point. An eigengene representation of all (C) upregulated and (D) downregulated DEGs in WNT (red) and NNT (blue). White/black bar indicates day/night period respectively. The red bar indicates the time period when WNT plants were exposed to higher temperatures. Error bars indicate ±SE (n=4).

The time of sampling greatly influences the identification of DEGs. The time point with the most DEGs was 1h before dawn, just before the WNT treatment ceased each day (396 DEGs, time point 23h). Only 12 DEGs were identified at dusk, the time point at which the WNT treatment was initiated. Even though the increased temperature was applied only from dusk until dawn, many genes were identified as DEGs in the samples taken during the day, when the conditions were identical between WNT and NNT. An eigengene representing the expression pattern of all genes induced at any time point by WNT highlights that the upregulated genes under WNT tend to show cyclic expression with a peak during the day in NNT. The timing of this peak in expression is altered by WNT (Fig. 2C). Functional enrichment of genes upregulated by WNT at any time point included MapMan^49^ terms enriched for protein posttranslational modification, signaling, carbohydrate metabolism, RNA processing, and kaurene synthesis (Fig. S4A). However, each time point presents a unique DEG profile and enriched functional categories (Supplemental Table 3). In part, this appears to be due to the underlying variation in expression in the control samples. For example, during the morning hours when photosynthetic genes are active, DEGs were enriched for photosynthesis-related activity, while at night no difference in expression was detected, likely due to their low value in control conditions at those time points. Therefore, sampling at only one time point would miss the impacts of WNT on molecular functions not active at that time. In fact, most of the 695 downregulated genes are identified during the nighttime. An eigengene representing the overall downregulated cohort indicates that under NNT, the majority of these transcripts peak just before dawn. In WNT, these genes have an overall lower amplitude of expression and an advance in the phase of peak expression (Fig. 2D). Downregulated genes were enriched for protein folding, photosynthesis, and heat stress (Fig. S4B).

### DEGs ARE ENRICHED FOR RYTHMICALLY EXPRESSED AND CIRCADIAN-CONTROLLED GENES

The observation that many DEGs show an overall change in the pattern of expression throughout the day (e.g., Fig. 2), and our hypothesis that the WNT disrupt the circadian clock, led us to speculate that the rhythmically expressed and circadian-regulated genes may have enhanced sensitivity to WNT. We evaluated if the DEGs in WNT are enriched for rice genes identified as rhythmically expressed in a previous study in controlled conditions by Filichkin et al.^44^. we observed significant overlap between transcripts we identified as DEGs in WNT and genes identified as rhythmic when grown in both photocycles and thermocycles (Fig. 3A). DEGs in WNT are under-represented for non-cycling genes in photocycles and thermocycles (*p*-value < 2.93e^-30^). The WNT DEGs were enriched for genes with a peak expression at night, between Zeitgeber times (**ZT**) 12-21h (Fig. 3A). Genes with peak expression at ZT19 (6h after dusk) showed the strongest enrichment for DEGs in WNT, (*p*-value < 8.55e^-5^). For example, *LOC_Os10g41550* is beta-amylase with rhythmic expression peaking just before dawn in the chamber-grown Nipponbare rice seedlings (Fig. 3B). We observe a similar peak in expression just before dawn in the corresponding gene, *MH10t0431700,* in our field-grown IR64 panicles in NNT. However, in WNT, the expression pattern is delayed, with a peak expression at dawn in WNT, when expression levels are already decreasing in NNT. The 24h expression pattern of beta-amylase in seedlings grown in photocycles and thermocycles has a higher correlation to NNT (0.95) than WNT (0.62). WNT DEGs are also overrepresented in transcripts rhythmically expressed in seedlings grown in only photocycles (Fig. 3C) or thermocycles (Fig. 3D).

**Figure 3.**
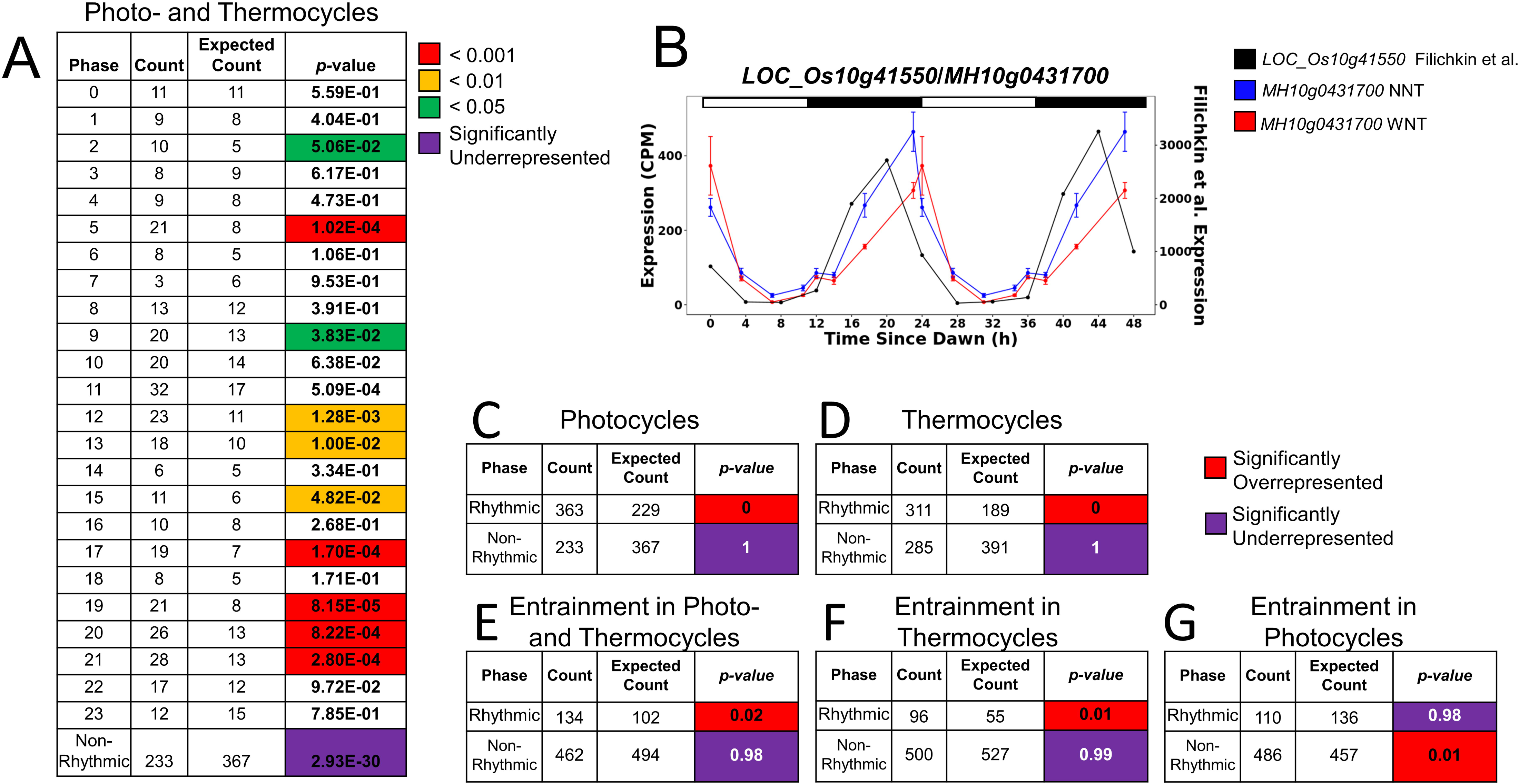
Rhythmic and circadian-regulated transcripts have increased sensitivity to WNT. Comparison of Warm Nighttime Temperatures (WNT) DEGs to a prior experiment examining diel rhythmic and circadian regulated expression of rice transcripts ^44^. (A) The enrichment of WNT DEGS compared to the phase of peak expression for rice transcripts when grown in photocycles and thermocycles. Enrichment is colored by *p*-value < 0.001 (red), <0.01 (orange), <0.05 (green), and underrepresented (purple). (B) Expression pattern of example transcript, *LOC_Os10g41550* (MH10g0431700) in conditions with both photocycles and thermocycles (black), Normal Nighttime Temperatures (NNT, blue), WNT (red). Enrichment of WNT DEGs in sets of transcripts identified as rhythmic or non-rhythmic from plants grown in (C) photocycles only, (D) thermocycles only, and (E) after entrainment in photo- and thermocycles. Enrichment of WNT DEGs in sets of transcripts identified as circadian-regulated after entrainment in (F) both photocycles and thermocycles, (G) thermocycles only, and (H) photocycles only. Significantly overrepresented (red) significantly underrepresented (purple).

In addition to rhythmic expression in the presence of photocycles or thermocycles, the WNT DEGs were also enriched for circadian-regulated genes with thermocycle-entrained expression. We considered genes to be circadian-regulated if the rhythmic expression persisted in constant conditions after entrainment in Filichkin et al. ^44^. The genes identified as DEGs in WNT showed enrichment for circadian-regulation only when compared to transcripts entrained in the presence of thermocycles. After entrainment with either thermocycles alone, or with both photocycles and thermocycles, WNT DEGs were enriched in the genes that maintained a rhythmic expression pattern when released to constant conditions (*p*-value < 0.05) (Fig. 3E, F). However, WNT DEGs were not enriched for in the transcripts that showed circadian regulation after entrainment with photocycles alone (Fig. 3G) indicating that WNT DEGs are enriched for genes under control of the thermocycle clock based on the data in Filichkin et al.^44^. For example, *LOC_Os10g41550* (the beta-amylase in Fig. 3B) is rhythmically expressed when entrained by thermocycles alone, but not by photocycles alone.

### WNT ALTERS TEMPORAL EXPRESSION PATTERNS

The enrichment of the WNT DEGs for genes previously identified as rhythmic in diel and circadian conditions suggests that WNT potentially disrupts the overall rhythmic expression of transcripts throughout the day. Therefore, we evaluated the rhythmicity of expression for all 15,213 detectible transcripts, even those not identified as DEG, in NNT and WNT grown samples. All transcripts were categorized as rhythmic or not rhythmic in these diel conditions using JTK cycle^50^ (*q-*value of <6.61e-15, Fig. 4A). The period of rhythmically expressed genes was similar in both NNT and WNT (Fig. S5).

**Figure 4.**
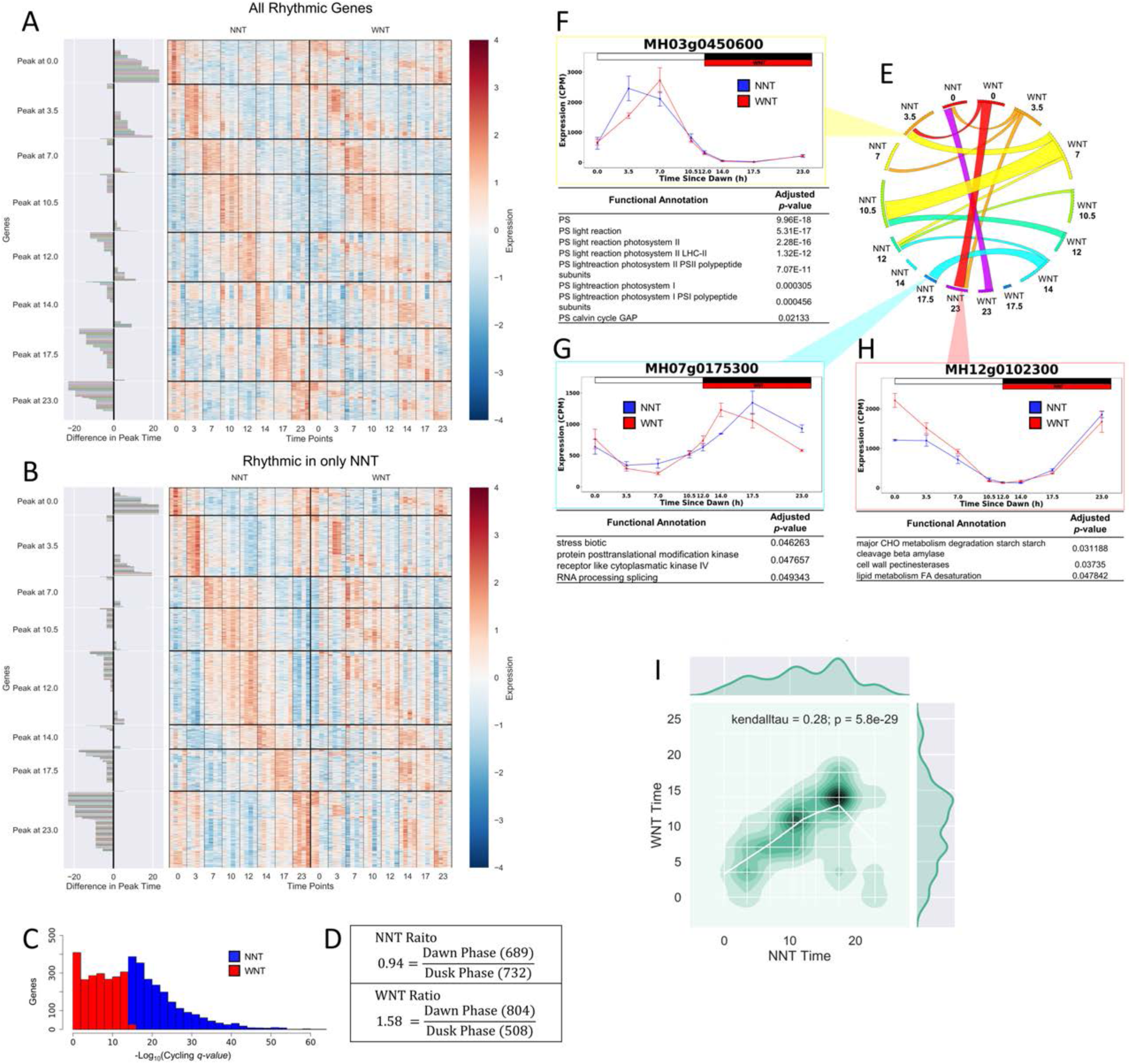
Patterns of expression are disrupted by WNT. (A) Heatmap of gene expression of rhythmic genes identified by JTK cycle in Normal Nighttime Temperatures (NNT) and Warm Nighttime Temperatures (WNT). Genes are ordered by NNT dynamics. The difference in peak time is the difference between the time of max expression in NNT compared to WNT. (B) Heatmap of gene expression of genes that are rhythmic in only NNT. The difference in peak time is the difference between the time of max expression in NNT compared to WNT. (C) -log_10_ (Q-*value*) distribution of NNT (blue) and WNT (red) JTK analysis of gene that cycle uniquely in NNT. (D) WNT alters the ratio of morning to evening peak phases of gene expression. (E) Circos plot of genes that are rhythmic in both NNT and WNT that show a change in the timing of their peak expression. Phase of peak expression in NNT is anchored on the left half of the plot, and phase of peak expression in HNT is on the right. Only transcripts which shown a change in peak expression are plotted. (F) Expression pattern of MH03g0450600 a transcript representative of genes with a peak of expression at 3.5h in NNT (Blue) that shifts to 7h in WNT (Red). Enrichment for MapMan ontology of genes with this 3.5h to 7h shift in peak expression are shown. G) Expression pattern of MH07g0175300 a representative transcript of the group of genes with a peak in expression at 17.5h in NNT (Blue) that advances to 14h in WNT (Red). Functional enrichment from MapMan Ontology is shown for genes with this shift (light blue). (H) Expression pattern of MH12g0102300, selected to represent the group of transcripts that peak at 23h in NNT and the peak expression is delayed until dawn in WNT. Function enrichment from MapMan ontology is shown for genes with this shift (red). (I) Density plot showing changes in timing of peak expression between NNT and WNT for genes identified as WNT DEGs.

We identified genes that were uniquely rhythmic in either NNT or WNT. Of the 6248 genes that cycled in NNT, 2136 genes lost rhythmicity in WNT (34%, Fig. 4B, Supplemental Table 4). The average *q*-value is 1.57e-16 in NNT; indicating that the genes that are losing rhythmicity in WNT were confidently identified as rhythmic in NNT (Fig. 4C). Genes that lost rhythmicity in WNT were enriched for heat response, protein folding, and amino acid metabolism (Supplemental Table 5). There were 1673 genes that were rhythmic only in WNT conditions (Supplemental Table 4). These genes that gained rhythmicity in WNT were enriched for DNA synthesis and chromatin structure, protein posttranslational modification, amino acid metabolism, cell wall synthesis, secondary metabolism, fatty acid metabolism, and cell cycle (Supplemental Table 5).

### WNT ALTERS THE PATTERN OF EXPRESSION OF GENES CLASSIFIED AS RHYTHMIC IN BOTH CONDITIONS

Even for genes that maintain a rhythmic pattern of expression in both NNT and WNT (66%, 4112 genes), the phase of expression may differ between the two conditions. As previously reported for chamber-grown plants, we observe a bimodal distribution of genes with peak primarily at dawn or dusk in NNT conditions (Fig. 4A)^43, 44, 51^. We classified genes as either morning phased (peaks at Dawn,) or evening phased (peaks at Dusk). In NNT conditions, the distribution between morning and evening phase was relatively equal (ratio of morning to evening phased genes = 0.94, Fig. 4D). However, in WNT this distribution is disrupted and there is an increase in the ratio of morning to evening phased genes (ratio = 1.58, Fig. 4D). The time of the maximum expression differs between WNT and NNT for 16.5% of the genes that maintain a rhythmic expression pattern in both conditions (Fig. 4E-H). WNT resulted in both delayed (Fig. 4F, H) and advanced (Fig. 4G) expression. This shift in peak expression is observed more often in select time points. More than 50% of the genes that in NNT peak at Dawn, 10.5h, 12h, 17.5h or 23h, have a shifted peak of expression in WNT. For example, *MH03g0450600,* a chlorophyll a/b binding protein, peaks in expression in NNT at 3.5h after dawn, but is delayed to 7h in WNT (Fig. 4F). Functional enrichment of all the genes with a similar delay in peak expression from 3.5h in NNT to 7h in WNT shows that this change in phase affects components of the photosynthetic machinery (Fig. 4F). Genes that peak just prior to dawn at 23h show an advance in their peak of expression in WNT. *MH07g0175300,* also known as OsNramp5, a metal transporter associated with disease resistance, peaks in expression at 17.5h in NNT but at 14h in WNT (Fig. 4G). This group of genes is functionally enriched with processes involved in biotic stress, protein phosphorylation, and RNA splicing (Fig. 4G). The expression of *MH12g0102300*, a GDP-L-galactose phosphorylase, peaks at 23h in NNT and plateaus in the early daytime hours. In WNT higher expression of *MH12g0102300* persists into the daytime hours changing the peak timing of expression (Fig. 4H). Genes with a similar disruption that delayed peak expression from 23h to Dawn in WNT were enriched for carbohydrate and fatty acid metabolism (Fig. 4H).

Many of the WNT DEGs showed a shift in their peak phase of expression (Fig. 4I). Genes that peaked during the daytime hours in NNT showed a smaller change in the timing of the peak expression compared to genes that peak after dusk in NNT, suggesting that the DEG calls may be due to the change in pattern of expression at night. Genes that peak in NNT after ZT12 showed a stronger effect, consistent with our previous observation of more DEGs post ZT12 (Fig. 2B), higher enrichment of rhythmically expressed transcripts in the nighttime period (Fig. 3A), and more genes that lose cycling after ZT12 (Fig. 4B).

### IDENTIFYING REGULATORS OF WNT PERTURBED TARGETS

To identify transcriptional regulators of WNT DEGs, we used two approaches in parallel (Fig. 5A). First we independently constructed a regulatory network from publicly available transcriptome data (External Data Network^52^). We constructed a second network only from our time course data (Internal Data Network). For the external data source we selected 555 Nipponbare microarray samples from Nagano et al.^52^, a dataset of field-grown rice samples with an emphasis on the time of day variation. Using sequential samples, we employed ExRANGES^53^ and GENIE3^54^ to construct a gene regulatory network, identifying the candidate targets of 1196 transcription factors (TFs) (Supplemental Table 7)^52^.

**Figure 5.**
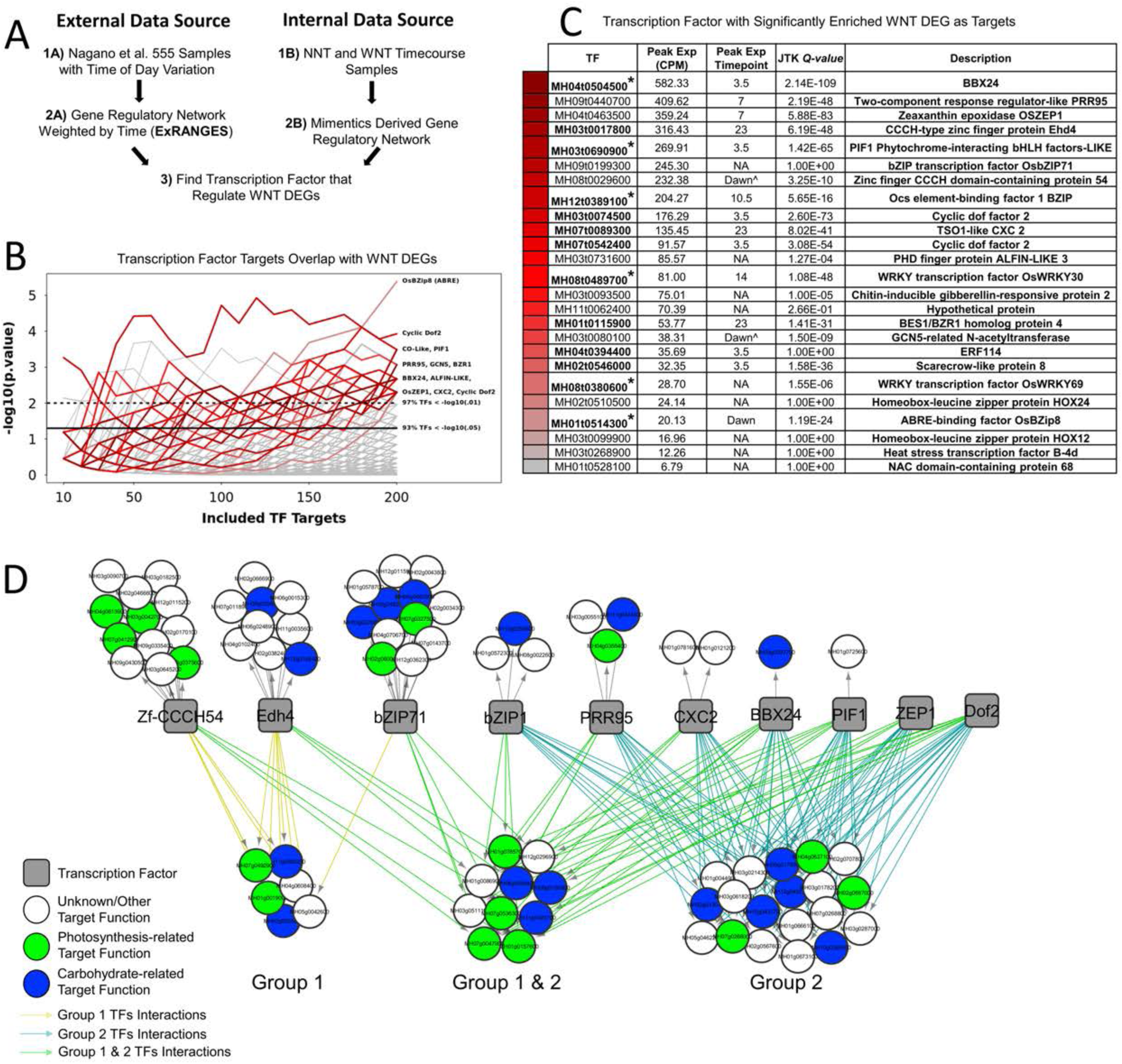
Inferring regulatory networks to identify WNT perturbed targets. (A) Workflow to establish Targets of Transcription Factors (TF)s. (B) Enrichment of TF targets for Warm Nighttime Temperature (WNT) DEGs from panicle time course. The enrichments score for the overlap of TF targets and DEGs is plotted on the y-axis as -log10(p.value). The enrichment test was performed for top 10-200 predicted TF targets. All annotated TFs were tested, each TF is represented by a grey line. Lines above dotted lines represent significance lower than a p.value of 0.01. Lines above dotted lines represent significance lower than a p.value of 0.05. TFs are colored by their peak expression (CPM) in the NNT time course. Red to green corresponds to high to low expression. (C) Table of significant regulators of WNT DEGs identified from external data source. TFs are sorted by their peak expression (CPM). * indicates significant regulators also identified by internal data source. (D) Overlap of the TF regulators and DEG targets. TF (grey), unknown/other targets (white), photosynthesis target (green), carbohydrate metabolism (blue) are connected by group 1 edges (yellow), group 2 edges (light blue), and group1/2 edges (light green). (E) Expression (CPM) of PIF1 at dawn (white) and dusk (grey) under night time temperatures of 24°C, 26°C, 28°C, and 30°C. All PIF1 targets and PIF1 targets that are also DEGs are separated into activated and repressed targets at dawn (white) and dusk (grey) timepoints. Line represents mean normalized expression (gene expression divided mean gene expression) of all targets at 24°C (green), 26°C (dark yellow), 28°C (orange), and 30°C (red). The shaded region represents standard deviation. (F) Expression (CPM) of PRR95 at dawn (white) and dusk (grey) under night time temperatures of 24°C, 26°C, 28°C, and 30°C. All PRR95 targets and PRR95 targets that are also DEGs are separated into activated and repressed targets at dawn (white) and dusk (grey) timepoints. Line represents mean normalized expression of all targets at 24°C (green), 26°C (darkyellow), 28°C (orange), and 30°C (red). The shaded region represents standard deviation between target gene expression.

Prior to evaluating the impacts of WNT on this network, we first established a confidence threshold for the TF target predictions and validated our network. We analyzed the overlap of the predicted TF targets with experimental Chip-Seq and binding motif data. Since few ChIP-Seq experiments in rice are available, we compared our identified targets with Arabidopsis ChIP-Seq experiments. In Arabidopsis, AtPIF4 *At*PIF4^55^ and *At*CCA1^56^ Chip-Seq data are available and prior work indicates that these Arabidopsis genes may be the functional orthologs of *Os*PIF1 and *Os*LHY respectively^57, 58^. We used orthologs identified between the rice variety MH63 and Arabidopsis^59^ to perform this cross-species comparison. We examined the enrichment of orthologs of ChIP-Seq targets for both Arabidopsis TFs to the predicted targets for the homologs of these two TFs in our rice network. The 500bp upstream promoter region of the *Os*PIF1 predicted targets is enriched for the cis-regulatory motif CACGTG, which is consistent with the known *At*PIF4 binding site (HOMER’s binomial cumulative distribution^60^ test *p*-value < 1E^-4^). Additionally, 26 of the 110 predicted targets for *Os*PIF1 overlapped with the Arabidopsis *At*PIF4 ChIP-Seq targets (*p*-value < 0.05). The predicted targets of *Os*LHY-chr8, were enriched for AAATATCT, which is also consistent with the known evening element binding site for *At*CCA1 (HOMER’s binomial cumulative distribution^60^ test *p*-value < 1E^-7^). We found that 24 out of the 108 predicted targets overlapped with *At*CCA1 ChIP-Seq targets (*p*-value < 0.05). The enrichment of our network for known targets and cis-regulatory motifs even across species, gives us confidence in the targets identified for each TF.

We identified the TF regulators whose targets, predicted from this external data set, had disrupted expression under WNT. TFs with targets that were enriched for WNT DEGs were considered regulators of WNT sensitive genes. The targets of 25 TFs were enriched for WNT DEGs (*p*-value cutoff < 0.01) (Fig. 5B). Most of these predicted regulators of WNT sensitive targets had a strong cyclic pattern of expression (JTK *Q-*value < 6E^-15^) and many were related circadian clock components (Fig. 5C). Of the TF regulators with high expression in panicle tissue, 35 target genes were shared between at least two TFs and 43 target genes have only one TF. Many of the target genes are related to photosynthesis and carbohydrate metabolism (Fig. 5D). Thirty of the 78 targets of these predicted regulators of WNT sensitive transcripts continue to cycle in constant conditions after entrainment in thermocycles alone (PHASER^56^ *p*-value < 0.01) indicating the target genes are enriched for thermocycle-entrained genes. A distinct pattern of TF and target grouping was apparent for the targets predicted to be regulated by more than one of these top candidate TFs (Fig. 5D). Group 1 consisted of targets regulated by Zf-CCH54, EDH4, and bZIP71, Group 2 consisted of targets regulated by bZIP1, PRR95, CXC2, BBX24, PIF1, ZEP1, and DOF2, and Group 3 targets had regulators from both groups. Homologs of many of the Group 2 regulators and targets have been previously associated with the circadian clock in Arabidopsis. For example, *MH03g0287000*, a Group 2 target which has sequence similarity to LNK1 in Arabidopsis, a known thermocycle-regulated clock component^61^. Of note, the Arabidopsis homolog of *Os*PIF1, *At*PIF3, has been previously identified as thermo-responsive^63, 64^. *At*PRR7 and *At*PRR9, the same family as *Os*PRR95, are regulators of temperature compensation^65^. BBX24 has been associated with circadian regulation^62–64^. Consistent with our observations of the altered rhythmicity of the transcripts themselves, the predicted regulators of the genes with expression altered by WNT suggest a link between the circadian clock and the observed altered expression under WNT.

### NETWORKS OF TRANSCRIPTIONAL RESPONSES ARE ALTERED IN WNT

Our Internal Data Network approach identified regulatory edges from the NNT and WNT expression data independently, based on^65^. This approach leverages the time series gene expression data and kinetic models of transcription regulation to estimate likelihood of TF regulation of other TFs (TF-TF). The results enable a direct comparison between the NNT and WNT constructed networks.

Of the 368 TFs that had regulators predicated with high confidence (TREP <= 3, BE<=0.25), the regulators of 89 of these TFs were perturbed by WNT. The perturbed network edges spanned the entire circadian cycle. TF-TF interaction of known circadian regulators also change based on the expression differences in NNT and WNT (Fig 6). For example, 17.5h after dawn four circadian TFs are expressed in NNT conditions, but in WNT there are no circadian TFs that peak at this time in the network (Fig. 6). Consistent with the External Network approach, BBX24 and PIF1 were identified as regulators with targets that have altered expression under WNT. However, regulators not present in the microarray-based Nagano et al.^52^ study can be detected in this Internal Data Network because the networks were generated directly from our Indica RNA-Seq data. For example, MH06g0689300, a Squamosa promoter binding protein-like (SPL) family member is identified in the Internal Data Network as a regulator of WNT DEGs, but is not on the rice microarray used by Nagano et al.^52^. Consistent with TFs identified as regulators of WNT DEGs from the External Data Network and the observed effects on patterns of gene expression, this analysis indicates that WNT may impact the functioning of the circadian clock in panicle tissue.

**Figure 6.**
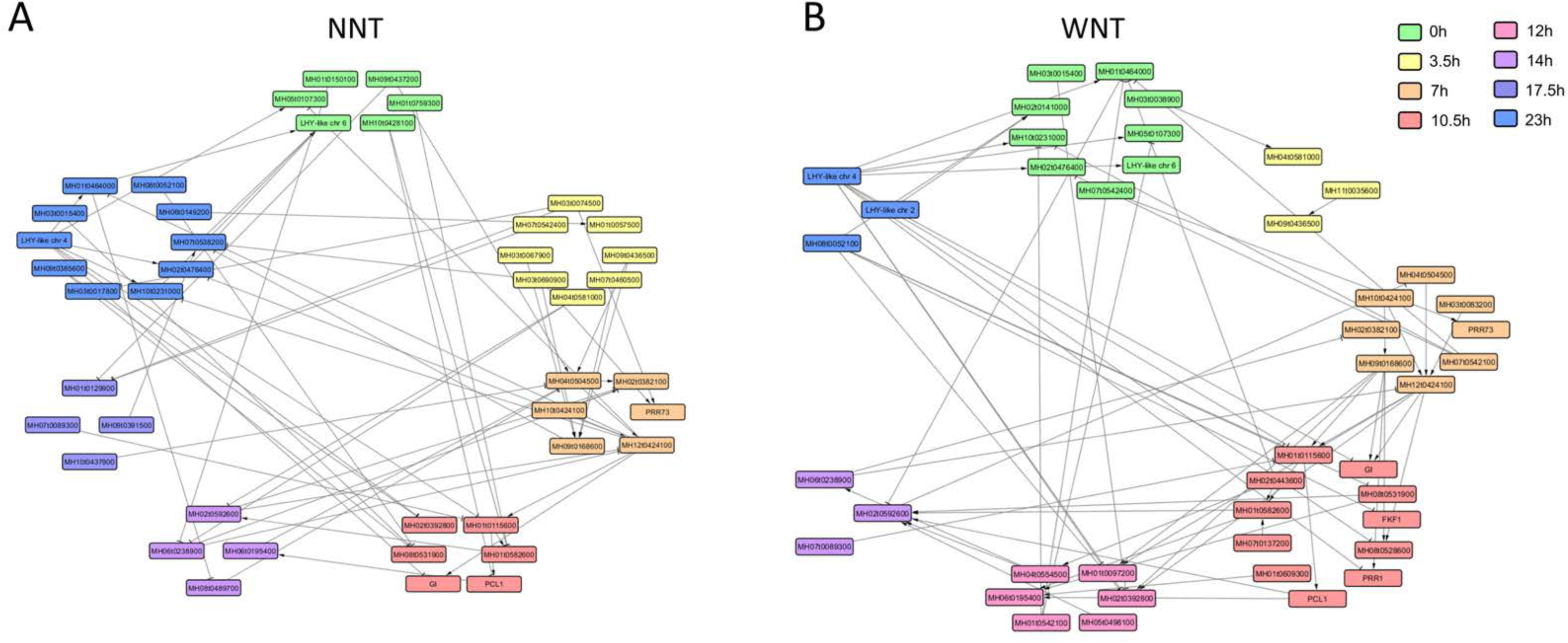
Distinct NNT and WNT networks from Internal Data. A network of Circadian/Circadian related TFs with TREP <= 3 and BE is <= 0.25 using A) NNT or B) WNT Internal Data. Nodes colored by peak expression (0h - Green, 3.5h - Yellow, 7h - Orange, 10.5h - Dark Pink, 12h - Pink, 14h - Purple, 17.5h - Dark Blue, 23h - Blue).

### VALIDATING TARGETS OF TFs THAT RESPOND TO INCREASING NIGHTTIME TEMPERATURE

Two independent network approaches identified overlapping regulators of WNT sensitive gene expression. Of the thirteen top regulators identified using the External Data Network that were also tested in the Internal Data Network, six were identified as regulators of WNT sensitive targets in both networks (Fig. 7A, Supplemental Table 8.)

**Figure 7.**
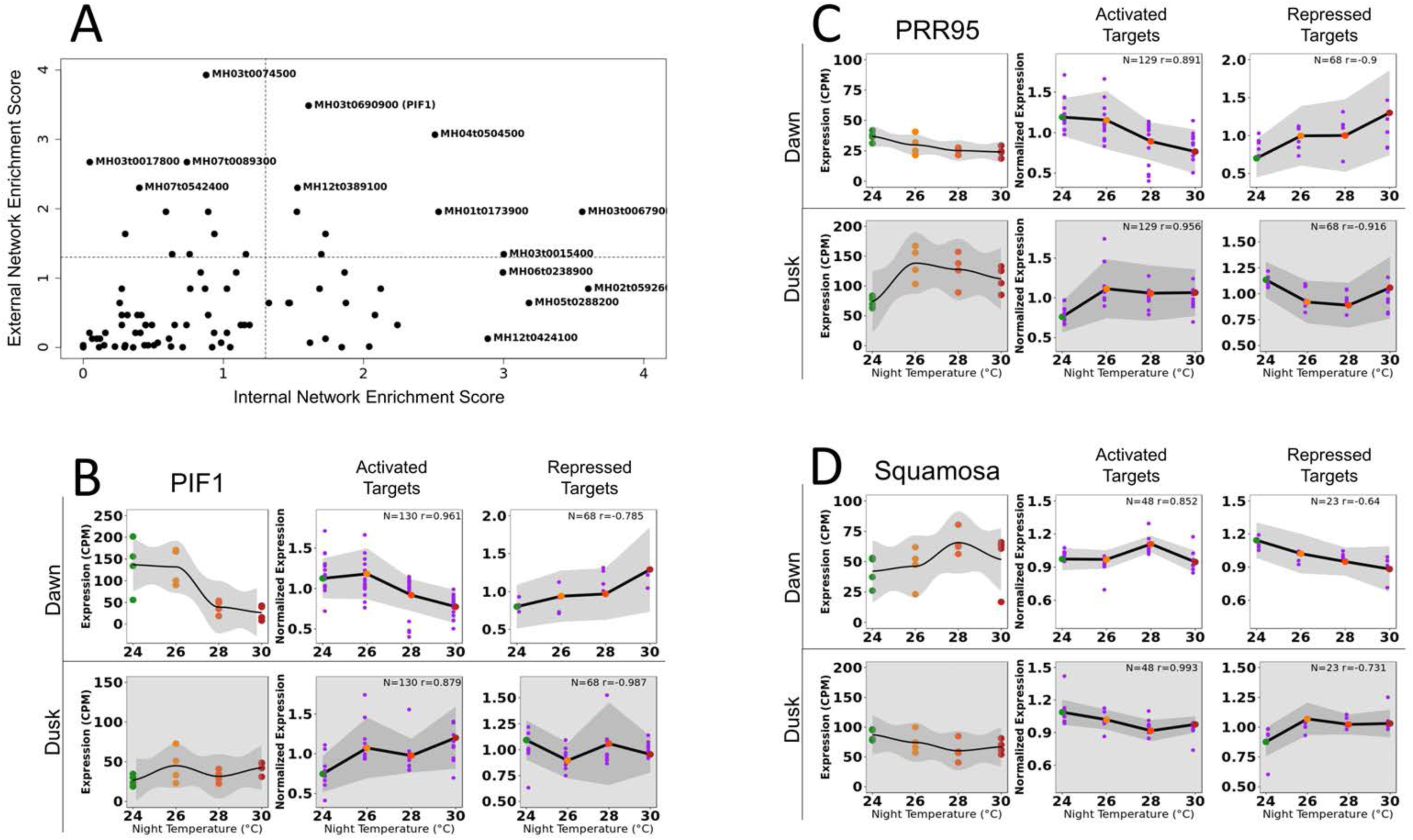
Evaluation of Regulators of Warm Nighttime Temperature Responses in an Independent Experiment. (A) Plot of enrichment of TF targets that are WNT sensitive using targets identified by the External (y-axis) and Internal network (x-axis). Enrichment Score is the - log2(p-value) of the hypergeometric test of the overlap between WNT DEGs and the predicted TF targets. Dotted lines are -log2(0.05) (B) Expression (CPM) of PIF1 at dawn (white) and dusk (grey) under night time temperatures of 24°C, 26°C, 28°C, and 30°C. All PIF1 targets and PIF1 targets that are also DEGs (purple) are separated into activated and repressed targets at dawn (white) and dusk (grey) timepoints. Line represents mean normalized expression (gene expression divided mean gene expression) of all targets at 24°C (green), 26°C (dark yellow), 28°C (orange), and 30°C (red). The shaded region represents standard deviation. (C) Expression (CPM) of PRR95 at dawn (white) and dusk (grey) under night time temperatures of 24°C, 26°C, 28°C, and 30°C. All PRR95 targets and PRR95 targets that are also DEGs (purple) are separated into activated and repressed targets at dawn (white) and dusk (grey) timepoints. Line represents mean normalized expression of all targets at 24°C (green), 26°C (darkyellow), 28°C (orange), and 30°C (red). The shaded region represents standard deviation between target gene expression. (D) Expression (CPM) of SQL at dawn (white) and dusk (grey) under night time temperatures of 24°C, 26°C, 28°C, and 30°C. All SQL targets and SQL targets that are also DEGs (purple) are separated into activated and repressed targets at dawn (white) and dusk (grey) timepoints. Line represents mean normalized expression of all targets at 24°C (green), 26°C (darkyellow), 28°C (orange), and 30°C (red). The shaded region represents standard deviation between target gene expression.

To evaluate the predicted regulators of WNT sensitive transcripts, we grew IR64 rice in field-based tents^13, 14^, where we could generate a gradient of night time temperatures (24°C, 26°C, 28°C or 30°C). We performed RNA-Seq from panicle tissue of plants grown in each of these night temperatures collected at the dawn and dusk time points. We examined the effect of this gradient of nighttime temperatures on the expression of the TFs and their predicted targets (Fig. 7B-D). Of the TFs predicted to regulate WNT DEGs, the expression of PIF1, PRR95, BBX24, SPL, and DOF2 TFs themselves responded to increasing nighttime temperatures (Fig 7, Fig. S6). For example, PIF1 expression is significantly reduced at dawn under 28°C and 30°C nighttime temperature conditions (Fig. 7B). At dusk, PIF1 is not expressed. Activated targets of PIF1 positively correlated with PIF1 expression at dawn (r = 0.96) with significantly reduced expression at 28°C and 30°C. Repressed targets of PIF1 negatively correlated with PIF1 expression (r = -0.79) at dawn and had significantly increased expression at 30°C. The correlation between PIF1 expression levels and the PIF1 targets predicted using the External Network that are also WNT DEGs is high for both activated targets (r = 0.97) and PIF1 repressed targets (r = -0.98) (Supplemental Table 9). In contrast to PIF1, PRR95 is only expressed at dusk and is differentially expressed at 26°C, 28°C, and 30°C (Fig. 7C). Targets predicted to be activated by PRR95 at dusk positively correlated with PRR95 expression (r = 0.96) and significantly increase expression in the increasing nighttime temperatures. At the dusk time point, targets predicted to be repressed by PRR95 correlated negatively with PRR95 expression (r = -0.92) and their expression was reduced under increasing nighttime temperatures. The predicted PRR95 Targets that are WNT DEGs are highly correlated with PRR95 expression at dusk (activated r = 0.94, repressed t = -0.87, Fig. 7C, Supplemental Table 2). The targets predicted to be activated by SPL, identified only in the Internal Data Network since it is not on the microarray, are also correlated with SPL expression both in dawn and dusk in the gradient experiment (dawn r= 0.85, dusk r= 0.99, Fig. 7D). The expression of the TFs predicted to regulate WNT sensitive targets showed > 0.7 absolute correlation value with their predicted targets, even though these targets were derived from a dataset not testing WNT^52^ (Figs. 5, 7, S6, Supplemental Table 8, https://go.ncsu.edu/clockworkviridi_wnt).

#### DISCUSSION

The asymmetric warming between day and night is an important environmental variable to consider for sustaining crop productivity in future climate scenarios. The physical phenomena of a significantly thinner planetary boundary layer during the night compared to the day, leads to larger effects on the surface air temperature during the nighttime^66^. WNT have a negative impact on large geographic regions, affecting agricultural productivity, compared to the more localized impact predicted for increasing daytime temperatures^7^. We observed the effects of WNT on field-grown rice plants. Prior studies have tested the effects of increased nighttime temperature in different genetic backgrounds, in chamber or greenhouse conditions, exposing plants to more extreme nighttime temperatures^16–18, 67^ that are greater than the nighttime temperatures predicted by IPCC or other climate models^1, 68^. However, a modest 2°C increase in nighttime temperature also reduced yield from 0-45.3% depending on year and variety^69^. From our treatment of similar temperature regime, we observed a significant 12.5% decrease in yield.

### WNT Alter the Timing of Fundamental Biochemical Processes

The term thermoperiodicity describes the differential impacts of day and night temperature changes on plants^70, 71^. Thermoperiodicity and the negative impacts of WNT have been observed in many crops and ornamental species^72–76^. Mechanisms for why a mild increase in nighttime temperature has such substantial impacts in contrast to a similar increase in daytime temperatures are not fully understood. Photosynthesis, transpiration, and respiration are temperature sensitive processes that contribute to yield. However, only respiration occurs consistently during the day and night. An increase in nighttime respiration has been associated with high nighttime temperatures^16, 17, 76–81^. Therefore, increases in dark respiration are often considered as the primary mechanism for the observed decrease in yield. However, the increase in maintenance, or nighttime respiration, may not fully explain the difference in yield^73, 82^. The impacts of high nighttime temperature during the vegetative stage can be compensated by active photosynthetic machinery^81^. In addition, the response of dark respiration under high nighttime temperature has only been documented in the leaf tissue either in rice^20, 81, 83^ or in wheat^84^, with no reports on panicle tissue.

Increasing nighttime temperatures have been associated with both positive and negative effects on the next day’s photosynthesis^80, 85–88^. The protective role of light-driven electron flow is not available in the dark, resulting in increased reactive oxygen species production and damage to photosystem II^89^. Therefore, the photosystems of many species of plants are more sensitive to heat-induced damage in the dark than in the light. Our findings that the early morning transcriptional induction of many of the photosynthetic genes is delayed in response to WNT may indicate that the altered timing of photosynthesis may compound the effects of increased respiration rates. Additionally, if the gene expression changes are reflective of the physiological timing, the delayed induction and subsequent overexpression later in the day we observe would result in variation in the observed difference in photosynthetic activity between control and WNT plants. Thus, the time of day the activity is measured could explain the conflicting reports of the effects of WNT on photosynthesis.

### The Phase-relationship as a Marker for WNT sensitivity

Examination of the transcriptional effects at a single timepoint would have missed potentially important transcriptional changes. For example, many of the upregulated genes are detected during the morning time points, while many of the downregulated genes are only downregulated during the nighttime hours. By examining transcripts at multiple time points, we can observe changes in the timing of expression that would be missed by a single time point. For example, we observe that many transcripts associated with photosynthesis show a delay in expression in the day and found changes in expression that would have been missed if the plants were sampled at the only one timepoint or were sampled only during the day. While any single time point captures at most 36% of the total WNT sensitive DEGs. However, **e**xamining the DEGs at Dawn and one hour before Dawn captures 59% of the total DEGs, and 79% of the total DEGs is captured by three time points (Dawn, 7h, and 23h).

### Disruption of Thermocycles may Contribute to the Negative Effects of WNT

While WNT can increase the rate of the biochemical activities occurring at night, another aspect of WNT is that they reduce the daily temperature range (DIF). This day to night temperature amplitude or thermocycle is beneficial to overall crop productivity^90^. Many crops have reduced yield under constant light conditions^91^, yet the damage caused by constant light can be reduced by providing a thermocycle with warm days and cooler nights in tomato and potato^92–94^. In tomato, net photosynthetic rates drop when grown in constant light and can be recovered by providing a recurring, daily temperature drop for just 2h, suggesting that the change in temperature acts as a cue to establish diel rhythms^95^. A linear relationship was observed between reduction in the DIF and adverse effects on morphology (e.g., reduced biomass) and physiology (e.g., increased nighttime respiration) in maize^76^. A large DIF could reduce the harmful effects of daytime heat on photosynthesis emphasizing the importance of the amplitude component of the daily temperature cycle. In Arabidopsis and rice, about 30-50% of the transcriptome is rhythmic under thermocycles in constant light, indicating that gene expression is one mechanism of responding to thermocycles^44, 96^.

In field conditions, we observe rhythmic expression of most transcripts, consistent with prior observations^97–99^. Our results in the rice panicle show that WNT globally disrupts the timing of gene expression in field conditions. Even after weeks of growth in increased nighttime temperatures, the transcriptional profiles differed between WNT and NNT panicles. Therefore, we propose an additional mechanism for the detrimental impacts of WNT. By disrupting the global patterns of gene expression, WNT disrupts the synchrony of molecular activities within the plant and the coordination with the external environment. The disrupted phase relationship between the timing of gene expression, downstream molecular events, and environmental factors results in reduced productivity. Although not well studied in rice, in Arabidopsis, plants that have a clock that is out of sync with their environment show decreased rates of growth and photosynthesis^39, 100, 101^. In barley, rhythmic expression of photosynthetic parameters showed allelic variation that suggests adaptation to local environments^102^. Therefore, the observed disruption of rhythmic expression could contribute to the overall decrease in yield and biomass (Fig. 1).

### Disruption of the Circadian Clock and Phytochrome Signaling by WNT

In Arabidopsis and rice, thermocycles can entrain the circadian clock and control the rhythmic expression pattern of up to 30% of the transcriptome^42–44^. In field conditions, we were not able to evaluate if the WNT effects persisted in constant light and temperatures. However, when we compared the WNT sensitive transcripts with prior data that identified circadian -regulated genes in rice, many of the WNT DEGs were circadian-regulated^44^. WNT DEGs are enriched for circadian-regulated transcripts when entrained by thermocycles alone or both light and temperature cycles in combination, but not when entrained by photocycles alone (Fig. 3). The enrichment for WNT DEGs in transcripts identified as thermocycle entrained suggests that WNT are disrupting the cues needed for proper timing of the expression of these transcripts. The transcripts rhythmically expressed in response to thermocycles are largely distinct from those entrained by photocycles^43, 44^. The functional significance of thermocycle-entrained genes has not been established in any plant species. However, if the negative impacts of WNT are due in part to the altered expression of thermocycle-entrained genes, understanding the roles of thermocycles in plant signaling and responses will be a critical component to anticipating the impacts of asymmetric changes in temperature patterns ^66^.

How the DIF is perceived and integrated into the circadian clock is unknown. Our gene regulatory network analysis identified 26 TFs with targets disrupted by WNT. Many of these TFs are known circadian clock regulators, associated with the circadian clock, or are themselves expressed rhythmically. We identified PIF1, a bHLH and Phytochrome Interacting Factor as a candidate regulator of WNT sensitive transcripts. In Arabidopsis, the OsPIF1 homologs, *At*PIF4 and *At*PIF5 are regulators of thermoresponsive growth^103–105^ and interact with circadian clock components^106, 107^. *At*PIF4 upregulates the auxin pathway, activating growth in response to increasing temperatures^108, 109^. Under increasing temperatures, the expression of *At*PIF4 and *At*PIF5 is regulated by *At*PHYB. *At*PHYB interacts with AtELF3, a component of the clock’s evening complex^106, 110, 111^. Loss of function mutations in the evening complex mimics the *phybde* loss of function knockouts under warmer temperatures^112, 113^. These interactions indicate that the photoreceptor PHYB connects changes in ambient temperature to the circadian clock in Arabidopsis^45–47^. Although the connections between phytochromes and increasing temperature are not as well studied in rice, if WNT affect the rate of interconversion between active and inactive forms of PHYB in rice, this could disrupt the output from the circadian clock through PHYB. The observed disruption of rhythmic expression of thermocycle-entrained genes could be a consequence of the impact on this PHYB signaling pathway. In Arabidopsis, the higher-order phytochrome mutants show a disruption in the timing of metabolite accumulation. In the loss of function mutants of four of the five Arabidopsis phytochromes, the plants accumulate sugars and amino acids to a higher level during the day and mobilize the sugars faster at night. Thus the reduced biomass of the phytochrome mutants is due to altered timing of photosynthesis and growth^114^. WNT grown rice plants also show a significant reduction in biomass (Fig 1) and alterations in grain quality^115–119^ suggesting changes in sugar mobilization.

Recent research suggests that the plant circadian clock is dynamically plastic to changes in the environment, which contributes to maintaining carbon homeostasis^41^. Feedback from the metabolic status, in part through endogenous sugar levels, can dynamically adjust the circadian oscillator depending on when the altered metabolism is perceived. These dynamic responses may explain contrasting shifts in expression we observe. For example, the delay in morning expressed photosynthetic transcripts and the advance in expression of genes with nighttime peaks in expression (Fig. 4E-H) may reflect temporally-varying WNT induced changes in carbon use and mobilization.

We also identified PRR95 as a candidate regulator of WNT sensitive genes. PRR95, a member of the Pseudo Response Regulator (PRR) family, is a known circadian clock component in rice^58, 120^. In Arabidopsis expression of PRR family members responds to temperature changes and *At*PRR7 and *At*PRR9 are important for temperature compensation, the ability of the clock to maintain a similar period across a range of temperatures^121^. Here we find that WNT alters the expression of *PRR95* (Fig. 7) and the predicted PRR95 targets. In our nighttime temperature gradients, we observed a high correlation between PIF1 and PRR95 expression and the expression of their predicted targets. This co-expression across a decreasing DIF supports the predicted regulatory role of PIF1 and PRR95 on these WNT sensitive transcripts and the effects of the thermocycle amplitude potentially through the circadian clock or phytochrome signaling on gene expression.

In addition to PRR95 and PIF1, we also identified other candidate regulators of WNT DEGs associated with the circadian clock. BBX24 is a CONSTANS like transcription factor which in Arabidopsis is an activator of PIF activity^64^. EDH4, a regulator of flowering and is essential in the environmental coordination of flowering^122^. Our findings suggest that WNT disrupt the temporal expression of transcripts by affecting the circadian clock or circadian-related components. However, we cannot distinguish if the effects on grain yield and biomass are due to direct effects of reduced amplitude on the entrainment of the circadian clock, through the downstream effects of increased nighttime respiration, or through temperature effects on transcription, translation, or other biological activities necessary for proper temporal coordination. However, by identifying the upstream regulators of the transcripts affected by WNT we can evaluate what processes may trigger the changes in gene expression.

In conclusion, growth under WNT results in global reprogramming of the rhythmic expression of the panicle transcriptome and multiple indicators suggest that this is through altered regulation of the circadian clock or the circadian-regulated outputs. It is unclear if the observed effects are a direct effect of WNT on the circadian clock machinery or an indirect consequence of increased nighttime respiration or altered metabolite transport. However, our results suggest that altered timing is a vital phenotype to monitor in response to WNT. The candidate regulators and their WNT-sensitive targets identified here can be used as markers to monitor the response to WNT across environments and genotypes and to identify the mechanisms to improve tolerance to WNT in anticipation of future weather patterns.

## ACKNOWLEDGMENTS

This project was supported by the Agriculture and Food Research Initiative competitive grant #2015-67013-22814 of the USDA National Institute of Food and Agriculture and the USDA National Institute of Food and Agriculture project 1002035.

## MATERIALS AND METHODS

### Crop Husbandry

Rice (*O. sativa* cv. IR64) was grown at the IRRI, Philippines during the dry season of 2014 (14° 11° N, 121° 15° E, 21 MASL). Seeds were exposed to 50 °C for three days to break dormancy. Pre-germinated seeds were sown in seeding trays and grown in the greenhouse for 14 days. Seedlings were then transplanted in the field at a spacing of 0.2 m x 0.2 m with one seedling per hill. Basal fertilizer composed of nitrogen (45 kg ha^-1^ N as urea), phosphorus (30 kg ha^-1^ P as single superphosphate), potassium (40 kg ha^-1^ K as KCl), and zinc (5 kg ha^-1^ Zn as zinc sulfate heptahydrate) was applied a day before transplanting. Plots were top-dressed with additional nitrogen during mid-tillering (30 kg ha^-1^ N), panicle initiation (45 kg ha^-1^ N), and heading (30 kg ha^-1^ N). Plots were kept fully flooded until just before harvesting at physiological maturity. Appropriate insecticides were used to control pest and disease damage.

Validation experiments under a gradient of nighttime temperatures were grown during the dry season of 2015. IR64 plants were pre-germinated and transplanted as described above and grown in a field-based tent facility at the IRRI, Philippines. Standard crop management practices including fertilizer application and pest control were followed^13^ and were similar to the 2014 experiment.

### Stress Treatment

An infrared heating facility using ceramic heaters was used to impose high temperatures during nighttime (1800-0600h) starting at panicle initiation until physiological maturity. The system has been described in detail in Gaihre et al. 2014^123^. The heated plots, with six ceramic heaters each, were programmed to have a temperature of +3 °C relative to the ambient temperature recorded from the control/reference plots. A total of eight rings were used in the experiment, with four replications per treatment. Temperatures were recorded throughout the day at 5-min intervals and monitored on a daily basis to ensure that the target temperature was achieved.

The physiological effects of WNT were measured by comparing rice grown in WNT to NNT. An increase in nighttime temperature in the field was achieved by planting rice inside a ring of ceramic heaters and maintaining a 3°C (actual: 2.36°C, SD ± 0.23) increase in air temperature inside the ring from dusk till dawn relative to ambient conditions (Fig. S1). The treatment was started from panicle initiation and persisted through the flower development and grain filling until physiological maturity. Throughout the experimental period the plants inside the WNT ring received the same temperature as the control plants during the daytime periods, but were consistently 2-2.5°C warmer at night (Fig. S1). The average diel amplitude in temperature was 9.1°C for NNT plants and was 6.7°C for WNT grown plants. For the entire period of stress imposition, the plants experiencing WNT did not receive a sudden increase in temperature, but rather a reduced decline in temperature (See insert Fig. S1).

Validation experiments in the temperature gradients were performed in field-based tents with programmed temperature control (for details see ^13^) were used to impose different nighttime temperatures. In brief, plants were exposed to ambient conditions during the day and exposed to target temperatures at night (18:00 – 06:00), starting from panicle initiation until physiological maturity. Temperature and relative humidity inside the tents were monitored using HOBO sensors installed above the canopy height^13, 14^. Plants were exposed to a total of four different night temperature conditions. Set temperature ranged from 24 – 30 °C with 2 °C increments. Heaters were used to impose 24, 26, 28, and 30 °C treatments. Two replicate tents were randomly allotted for each of the treatments.

### Sample Collection

Whole panicles at the 50% flowering stage (i.e. upper 50% of the panicle has just finished flowering) were collected. Uniform sets of panicles were identified, tagged, and collected when the middle portion of the panicle had spikelets undergoing anthesis or flower opening (time point dawn + 3.5h) or had just flowered (all other time points). The tagged panicles were collected at dawn (6:15AM), dawn + 3.5h, dawn + 7h, dawn + 10.5h, dusk, dawn + 14h, dawn + 17.5h and just before the next dawn, i.e. dawn + 23h. For each treatment, four replicates per time point were collected. Samples were collected in tubes and immediately immersed in liquid nitrogen, after which were stored at -80 °C until sample processing.

For the nighttime temperature gradient samples, main tillers with panicles at 50% flowering were identified and tagged for sample collection. Panicles were collected immediately after sunrise and sunset. For each controlled nighttime temperature, two replicate panicles were collected per tent, yielding a total of four replicates per treatment. The 24°C nighttime temperature was closest to the average NNT conditions, while the average nighttime temperature of our WNT samples described above was closer to 26°C (Fig. S1). Samples were collected in 50 mL falcon tubes and immediately immersed in liquid nitrogen, and stored at -80°C until analysis.

### Agronomic Characterization

Twelve hills from each replicate plot were harvested at physiological maturity to measure yield and yield components including total aboveground biomass, number of spikelets per panicle, 1000-grain weight, panicle per m^-2^, and spikelet fertility (Supplemental Table 1). Samples were processed according to standard practices following Lawas et al.^124^. To evaluate the effects of WNT treatment on each trait we performed the Kruskal-Wallis Test using R version 3.4.1.

### Library Preparation

The upper 50% of the panicle was first ground in liquid nitrogen with a metal pestle. The tissue was then lyophilized at -60°C overnight before RNA extraction. Total RNA was extracted using the RNeasy Plant Mini Kit (Qiagen) with the recommended RLT lysis buffer. The provided RNA extraction protocol was followed by the inclusion of DNase treatment. After the RWI wash step, 3 uL of DNaseI (10U/ul Roche), 8 uL buffer (200 mM Tris, pH 8.0, 20 mM MgCl2, 500 mM KCl), and 69 uL nuclease-free water was added to each column and incubated at room temperature for ten minutes. Following DNase treatment, the column was rewashed with the RWI buffer from the Qiagen kit. The RNA concentration was then measured with NANOdrop 2000 (Thermo Scientific). Two micrograms of RNA were used with NEBNext Poly(A) Magnetic mRNA isolation kit (NEB). Oligo(dT) attached to magnetic beads isolate mRNA by attaching to ploy(A) modified mRNA. Before library preparation, mRNA was heated to 95°C for the recommended 15 minutes to achieve 150-200bp fragment sizes. NEBNext Ultra RNA Library Prep Kit for Illumina was then used to prep mRNA for sequencing. cDNA was prepared with random hexamers and a Protoscript II reverse transcriptase for the first strand synthesis and followed by second strand synthesis. DNA was size selected using AMPure beads (Beckman Coulter) after end repair and adaptor ligation to isolate 150-200bp fragments. PCR library enrichments was then done with the inclusion of USER step for strand specificity. Fifteen cycles (the maximum recommended number of cycles) was used which did not result in any over amplification peaks. To measure library quantity and quality, samples were analyzed on an Agilent Bioanalyzer high sensitivity DNA chip after a 1:4 (or 1:10 if the concentration is too high) dilution and quantified using the NEB library qRT quantification kit. Libraries were diluted to 10nmol/ul concentrations before sequencing. Sequencing was done at North Carolina State University’s Genomics Science Laboratory on the Illumina Hiseq 2000.

### RNA-Seq Data Alignment and Quantification

After FastQC, fastq files were trimmed by 10nts with seqtk Trim fastq. Option used: -b -10. Fastq files were trimmed by 10nts before being aligned. Trimmed files were aligned using tophat v2. Reads were mapped to the indica variety MH63 and mapped the reads to the MH63 reference genome^59, 125^. The reference genome and annotation files were obtained from http://rice.hzau.edu.cn/rice ^59^. Mismatches allowed was 2, read gap length was 2, library type was set to first-strand, and the rest of the parameters remain as the default. The resulting bam files were then piped to htseq count version 0.6.0. Using the options -f bam, -s reverse, and -m intersection-nonempty counts per gene were generated for the annotated genes in MH63. Each sample had 80% or greater reads mapped. We observed low variability between replicates within both NNT and WNT groups (Fig. S2 and S3).

### Differential Expression

The raw count matrix was used for differential expression analysis with EdgeR version 3.10.5 ^126^. The genes were first filtered by counts; genes with more than 10 counts in four samples were kept. Normalization factors and dispersion estimates were then done according to the EdgeR pipeline. Differentially expressed genes (DEGs) between plants exposed to WNT or NNT at each time point were identified (Fig. 2) using EdgeR’s generalized linear models and a likelihood ratio test determined differential expression of transcripts. A FDR < 0.05 & logFC > 0.5 was considered to be differentially expressed. R version 3.2.1. Filters were applied to remove transcripts with less than 10 counts in less than 4 samples. Using filtered and aligned reads, transcript levels were used to find differentially expressed genes at different times of day. Differentially expressed (DE) genes were identified using EdgeR, comparing each time point between HNT treated and control samples (FDR < 0.05 & logFC > 0.5).

### MapMan Enrichment

All gene ontology tests were done with MSU annotations obtained from http://www.gomapman.org/ontology ^49^. Enrichment was then calculated using all MH63 genes as the background and either DE genes or genes that lost rhythmicity in HNT as the input. Phyper was used to calculate the p-value. After *p*-value adjustment for multiple testing corrections, a term with <0.05 FDR and a minimum of five genes per functional term were considered to be enriched.

### Enrichment of DEG in Diurnal and Circadian datasets

PHASER ^127^ was used to look for enrichment of rhythmically expressed genes in 50% flowering time course ^127^. We limited our comparison to the 596 WNT DEGs that had orthologs to Nipponbare and were on the A-AFFY-126 (Affymetrix, Santa Clara, CA) used by Filichkin et al., ^44^. After conversion of MH63 IDs to Nipponbare MSU orthologs, all DEG genes were used as an input to Phaser. The LDHC and LLHH_LLHC conditions of *Oryza sativa* data were used as the background. A correlation cutoff of 0.7 was used.

### JTK Cycle

JTK cycle R script was taken from http://www.openwetware.org/wiki/HughesLab:JTK_Cycle ^50^. To stringently identify differentially cycling genes, circadian core genes^98^, were plotted in the NNT data. The transcripts were visually inspected, ranked by JTK *q*-value, and a cutoff of the JTK *q*-value was set for the last gene that was clearly rhythmically expressed (MH03t0197300, 6.617080673040279e-15 inclusive). This *q*-value cutoff resulted in 6248 periodic in NNT, 4112 periodic in NNT and WNT 2136 periodic in NNT but not WNT. Full statistical results are in Supplemental Table 10.

### Inference of Regulators in External Data Network

Transcription regulators for rice were identified by RiceSRTFDB^128^. From Nagano et al. we exported 555 samples and created 15 different time series sets^52^. The data include samples taken in intervals of every ten minutes, every two hours, and every 12 hours from different developmental stages and tissue types (Supplemental Table 6). The dataset was subset to RAP-IDs that had orthologs to MH63 IDs^59^. The microarray data were ordered by sample type to create a time sequential series for each sample type. Sample divisions are given in supplemental table 3, and ExRANGES was applied to a series that contained time information. ExRANGES weights gene expression by the significance of the change in expression and was used because it performs optimally on large time course datasets. ExRANGES parameters cycle was set to FALSE and the sample size was set to 1000. The weight expression output of ExRANGES was then used to construct a transcription factor regulatory network. To predict regulatory interaction between the transcription factor and the target gene, GENIE3 source code was downloaded from http://www.montefiore.ulg.ac.be/~huynh-thu/software.html54. For GENIE3, we used 1000 trees. The importance measure was then ranked for each TF and its targets. To validate network results, we used the top 200 predicted targets for PIF1 and LHY-Chr8. We compared ChIP-Seq targets for AtPIF4 and AtCCA1. Orthologs between RAP-IDs and Arabidopsis were converted using mappings downloaded from rigw.org^59^. To test for motif enrichments, Homer’s findMotif.pl (v4.10) function was used on an imported version of the MH63 genome^60^. The motif search was limited to 500bp upstream of the transcription start sites and 50bp downstream of TSS. The motif length was limited to 6, 8, and 10bps. To test for enrichment of DEGs, MH63 Ids were converted to RAP Nipponbare orthologs using rigw.org^59^. Using phyper, enrichment was calculated for the overrepresentation of WNT DEGs in TF targets. We tested enrichment for WNT DEGs for all TFs. We looked at enrichment in the top 10-200 predicted targets for each TF.

### Inference of Regulators in Internal Data Network

For comparison of gene regulatory networks from control and WNT panicle tissue, the edge finding pipeline from McGoff et al.^65^ was employed. The predicted regulators of each TF were determined by constructing a regulatory network consisting of TFs with homology to known circadian regulatory TFs and determining their potential for regulating the TFs with altered expression in WNT. The model considered the relationship between each potential regulator and the altered TFs of interest using the gene expression dynamics observed by RNA-Seq^65^. A model distribution was constructed to determine the topological differences between model gene regulatory networks in NNT and WNT. First, dynamic gene profiles were selected for analysis. Rice Transcription Factors were first analyzed by DEseq to determine if they were differentially expressed in Control and WNT samples (FDR < 0.05). Next, using the JTK periodicity finding algorithm, highly periodic TFs from control and WNT samples were identified, considering the possibility that a TF may be highly periodic in one dataset and not another. Also, TFs that demonstrated periodic dynamics in both control and WNT were added, even if their peak times were not consistent between experiments. Finally, known circadian regulators were added. All told, 368 genes were considered, of which 356 were Transcription Factors. Using the edge finding pipeline, only TFs were considered for regulators, while all 368 genes were target nodes.

From the edge finding algorithm, the following attributes of each possible regulator node to target node interaction were collected. Replicate 2 from control tissue and Replicate 2 from WNT were used for network analysis. Edge Probability (EP) – the likelihood that an edge exists between a given regulator and a given target. This measure considers the goodness-of-fit of the regulation simulation to the observed data and a number of other factors. The score ranges from 0 to 1 and the sum of all scores for a single target is 1. The higher likelihoods have larger values. Mean Squared Error (MSE) – the mean squared error for the regulation simulation compared to the observed data. This is always a positive number. Smaller values indicate a better fit. Baseline Error (BE) – compares the error for the simulation versus the error for a straight line fit. The output ranges from 0 to 1. Smaller values indicate a better fit. This provides a normalized goodness of fit measure. Target Rank Edge Probability (TREP) – shows how a given edge ranks against other edges leading to the same target. The smaller values indicate better rankings.

An inspection of the results led to construction of Control Panicle gene regulatory networks that satisfied the following criteria: The TREP must be in the top 3 potential regulatory edges for each target node. The BE must also be less than 0.25. Finally, each TF node must have at least one edge “in” and one edge “out” indicating that it is both a target and regulator of another TF in the network. These score cut-offs result in a gene regulatory network model that allows for comparison to the WNT Panicle GRN. To determine differences in network topology, regulator targets in WNT were identified when their regulator profile were altered (different regulators in the top 3 TREP and altered incoming edge scores). For each target, non-parametric correlation between regulator scores from control and HNT (Baseline Error) were computed. If there was no significant correlation (p > 0.05), this target is perturbed. 193 genes in the network had significant positively-correlated regulation between NNT and WNT. 89 genes in the GRN constructed in NNT were perturbed in the WNT panicle. Network visualizations using these criteria were constructed in Cytoscape^129^.

### Supplemental Figure Legends

**Supplemental Figure 1** Experimental setup of WNT field samples. (A) Image showing the growth of the WNT samples, contained within the ring of ceramic heaters that maintained (B) Actual temperature data from the period of panicle initiation, when the heaters were turned on for WNT plants until harvest. (C) Close up of four days of treatment showing that the day to night temperature difference in both NNT and WNT exceeds the nighttime temperature difference between NNT and WNT.

**Supplemental Figure 2** Violin plot showing variation between the replicates of samples from Normal Nighttime Temperatures (NNT). Pairwise comparisons of variation between each of the four NNT replicates. The distribution of the Kendall-Tau dissimilarity score (Score) for all transcripts is shown.

**Supplemental Figure 3** Violin plot showing variation between the replicates of samples from Warm Nighttime Temperatures (WNT). Pairwise comparison of variation between each of the four WNT replicates. The distribution of the Kendall-Tau dissimilarity score (Score) for all transcripts is shown.

**Supplemental Figure 4** Functional categorization and summarized expression of WNT DEGs. Mapman-based functional enrichment of genes (A) upregulated and (B) downregulated in response to Warm Nighttime Temperatures (WNT) compared to Normal Nighttime Temperatures (NNT).

**Supplemental Figure 5** The period expression of most genes is similar between Warm Nighttime Temperatures (WNT) and Normal Nighttime Temperatures (NNT). Density of genes with difference in period lengths indicated between WNT and NNT.

**Supplemental Figure 6.** Validating targets of other TFs responding to increasing nighttime temperature. Expression (CPM) of (A) Dof2, (B) BBX24, (C) bZIP71, (D) EDH4, (E) bZIP1, (F)? (G) Zf-CCCH54, (H)ZEP1, at dawn (white) and dusk (grey) under nighttime temperatures of 24 C (green), 26 C (dark yellow), 28 C (orange), and 30 C (red). Mean of (A) Dof2, (B) EDH4, (C) Zf-CCCH54, BBX24, bZIP1, ZEP1, bZIP71 targets (black) and Dof2, EDH4, Zf-CCCH54, BBX24, bZIP1, ZEP1, bZIP71 targets that are also DEGs (purple) are separated into activated and repressed targets at dawn (white) and dusk (grey) timepoints. The line represents mean normalized expression (gene expression divided mean gene expression) of all targets at 24 C, 26 C, 28 C, and 30 C. The shaded region represents the standard deviation.

**Supplemental Table 1** Agricultural Traits for IR64 plants grown in Warm Nighttime Temperatures (WNT) and Normal Nighttime Temperatures (NNT).

**Supplemental Table 2** Genes identified as DEG in Warm Nighttime Temperatures (WNT) compared to Normal Nighttime Temperatures (NNT) at each time point. Log fold change (LFC) and FDR corrected significance (FDR) are shown. Positive LFC indicates the transcript is upregulated in WNT.

**Supplemental Table 3** Functional enrichment of genes identified as DEG in Warm Nighttime Temperatures (WNT) compared to Normal Nighttime Temperatures (NNT) at each time point using MapMan 49 terms.

**Supplemental Table 4** Genes identified as cycling in Normal Nighttime Temperatures (NNT), in Warm Nighttime Temperatures (WNT), cycling only in NNT, and cycling only in WNT.

**Supplemental Table 5** Functional enrichment of genes identified as rhythmic either only in Normal Nighttime Temperatures (NNT) or only in Warm Nighttime Temperatures (WNT).

**Supplemental Table 6** Samples used to generate External Data Network. The 555 samples from ^52^ that were used to construct the External Data Network. These were grouped into 15 series and the table indicates if ExRANGES was applied to each series.

**Supplemental Table 7** The gene identifiers of the transcription factors considered as regulators in ExRANGES for generation of External Data Network.

**Supplemental Table 8** Enrichment of the predicted targets of each regulatory transcription factor (TF) for differentially expressed genes (DEGs) under warm nighttime temperatures (WNT). Each TF tested for the External Data Network (1174 TFs) and the Internal Data Network (356 TFs). The enrichment score is the -log_10_ (*p*-value) of the predicted targets of that regulator for WNT DEGs. The six TFs enriched in both networks are indicated.

**Supplemental Table 9** Correlation between top candidate regulatory transcription factors (TFs) and their predicted targets in the validation assay using a gradient of warmer nighttime temperatures (WNT). The correlation between the top candidate TFs and all the predicted targets (All Targets) or the targets that were identified as WNT DEGs (WNT DEG Targets) in the WNT gradient experiment.

