## Supplementary figures and images for "Warm nights disrupt global transcriptional rhythms in field-grown rice panicles"

### Figure S2

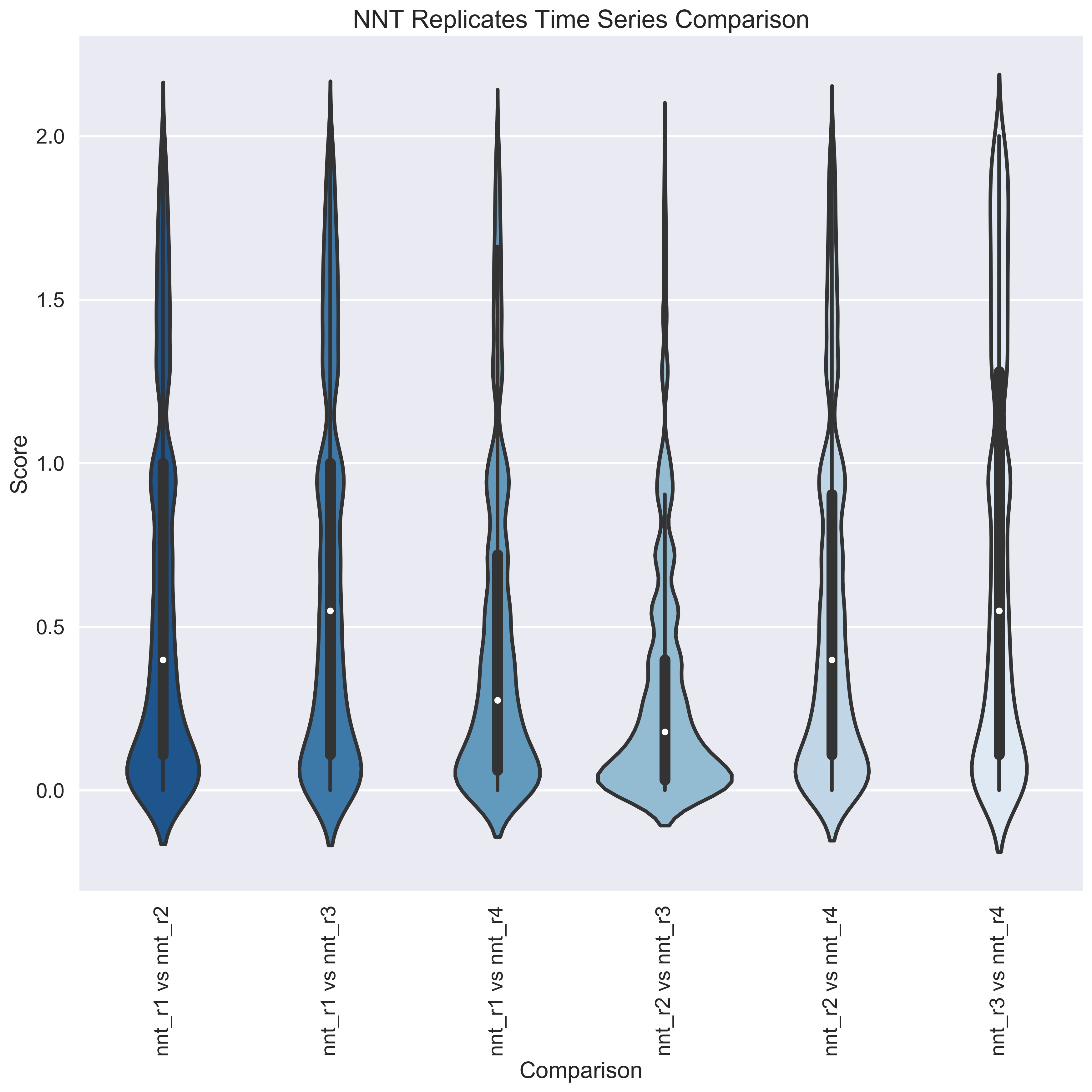

### Figure S3

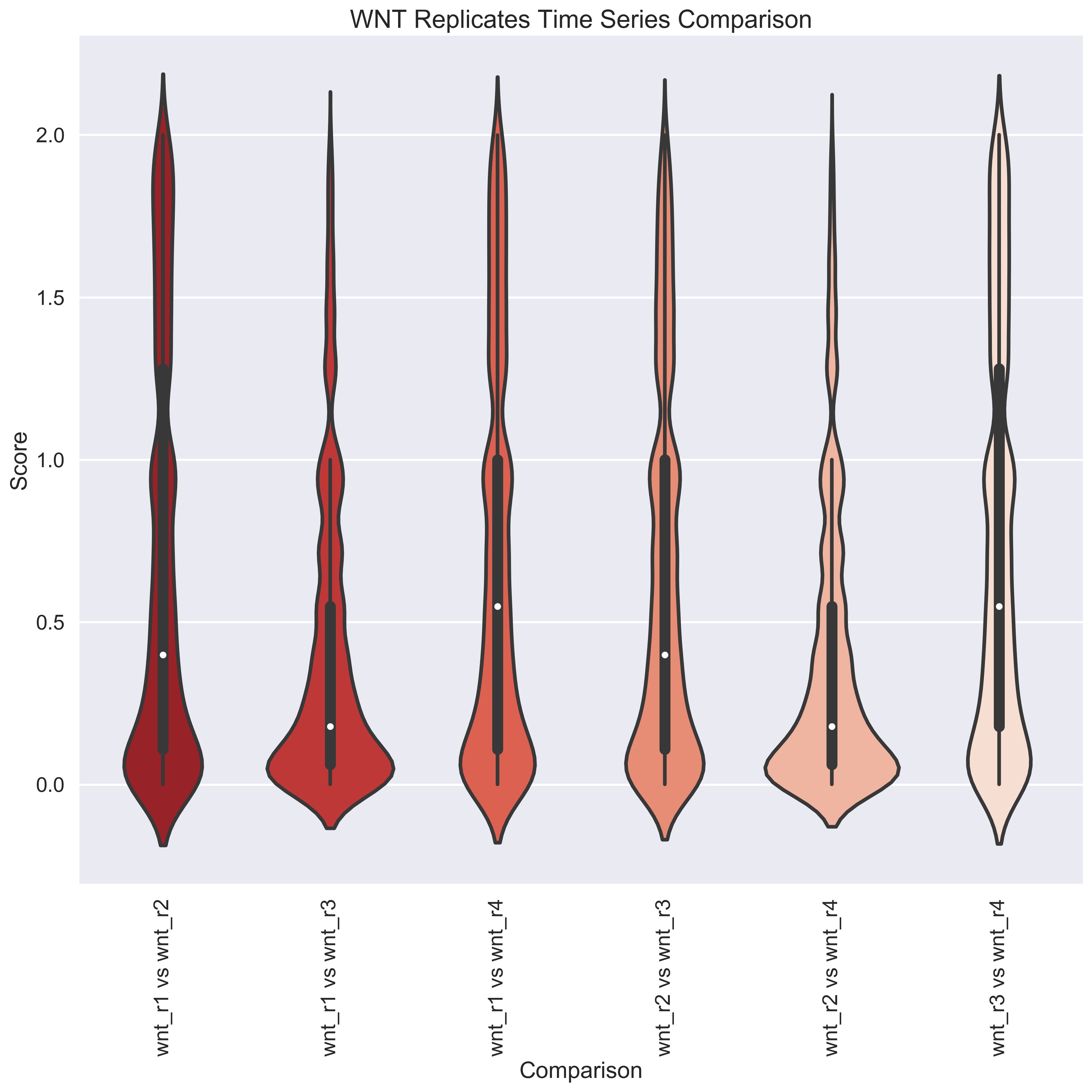

### Figure S3

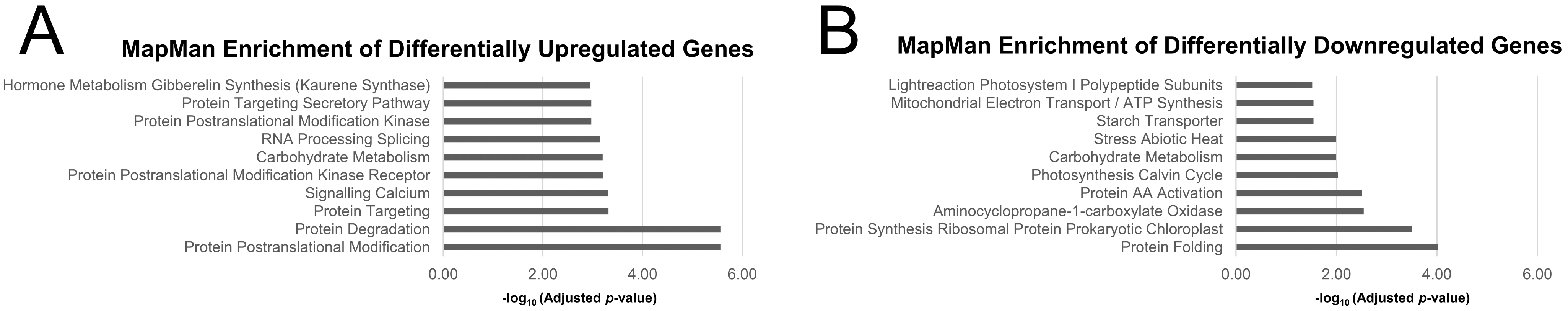

### Figure S5

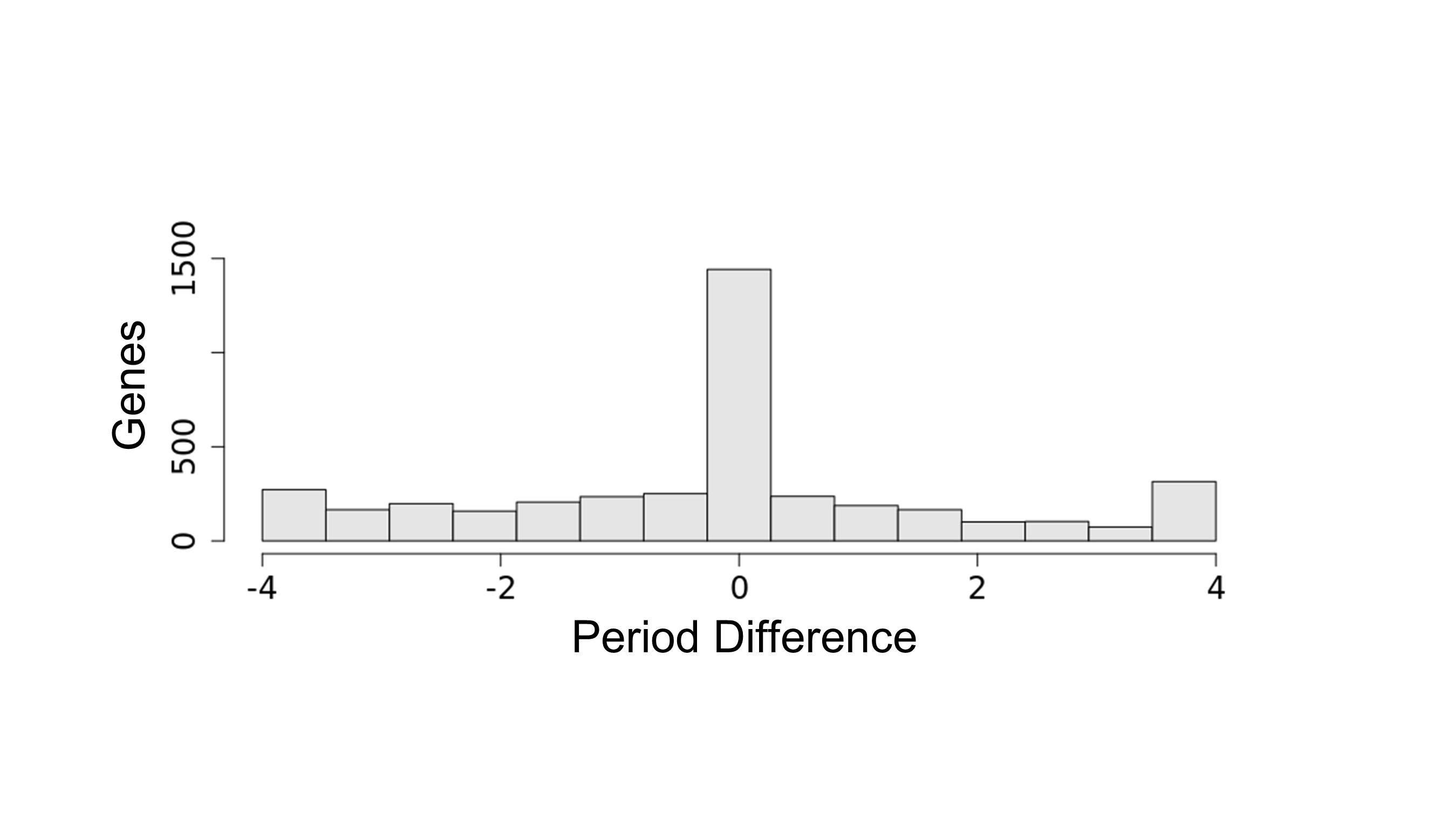

### Figure S6

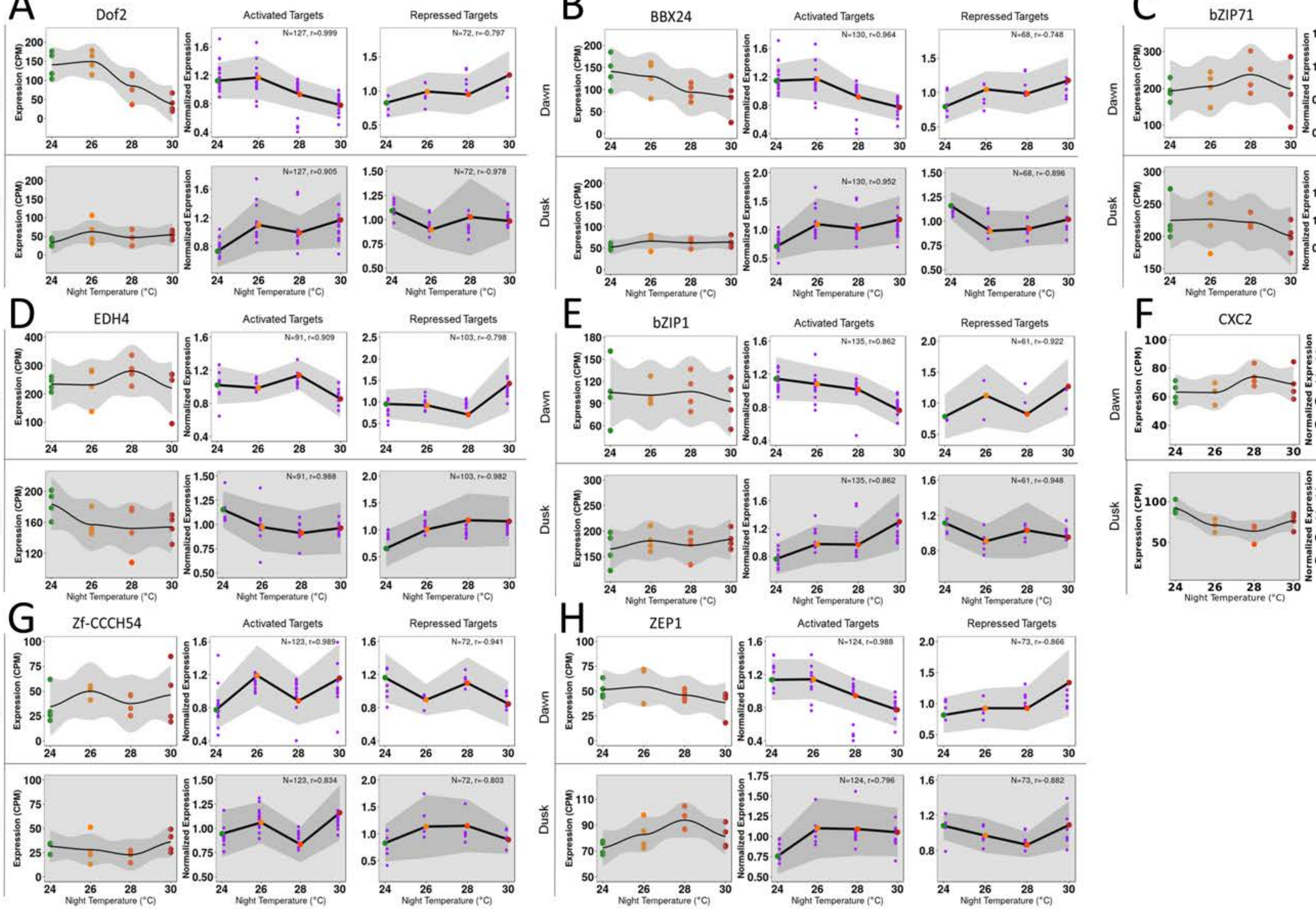
